# Proton channel inactivation results in loss of chloroplast NDH complex activity

**DOI:** 10.1101/2024.10.01.616091

**Authors:** Deserah D. Strand, Stephanie Ruf, Omar A. Sandoval-Ibáñez, Ralph Bock

**Affiliations:** Max-Planck-Institut für Molekulare Pflanzenphysiologie, Am Mühlenberg 1, D–14476 Potsdam-Golm, Germany

**Keywords:** Photosynthesis, proton transport, site-directed mutagenesis, chloroplast, bioenergetics

## Abstract

The plastidal photosynthetic complex I (formerly NAD(P)H dehydrogenase-like complex, NDH) remains enigmatic in its function within the electron transport chain of higher plants. While the NDH complex shares high homology with complex I, a key component of the respiratory electron transport chain, electron transport rates through the NDH complex in thylakoids are relatively low. In this study, we took a structure-function approach and mutated the plastid genome-encoded *ndhF* gene to abolish the NdhF proton channel of the NDH complex. These mutations led to loss of plastoquinone reductase activity, indicating tight coupling between the proton and electron transfer reactions within NDH. Additionally, loss of the transverse helix of NdhF led to loss of the NDH complex, suggesting that this region of the NdhF subunit is required for complex stability. In agreement with previous studies using *ndh* knockout mutants, loss of NDH complex activity did not result in measurable changes in rates of steady-state cyclic electron flow. However, all mutants displayed a shift in the sensitivity of pH-dependent feedback regulation of the photosystem II antennae to total protonmotive force (*pmf*), indicating a defect in either stromal redox state or *pmf* distribution into ΔpH and Δψ.

## Introduction

Photosynthesis has multiple regulatory points to maintain efficiency while avoiding photodamage. One of these regulatory points is the reduction of the plastoquinone (PQ) pool by stromal reducing equivalents (e.g., ferredoxin, Fd; or NADPH). This reduction of the PQ pool from the stroma and subsequent plastoquinol (PQH_2_) oxidation at the cytochrome *b*_6_*f* (*bf*) complex, termed cyclic electron flow (CEF) around photosystem I (PSI), bypasses photosystem II (PSII) to increase proton translocation and ATP production per reducing equivalents generated at PSI, to supply additional ATP to the chloroplast. Additionally, CEF proton translocation into the lumen adds to protonmotive force (*pmf*), of which the ΔpH component is regulatory, thus activating the downregulation of the PSII antennae (energy-dependent quenching, *q*_E_) and slowing electron transport through the *bf* complex (photosynthetic control, PCON) [reviewed in (*1*)]. There have been several routes of CEF identified in plants and algae, the most efficient route of which is the photosynthetic complex I (formerly NADPH dehydrogenase-like complex, NDH) (*2–5*).

The NDH complex is homologous to the respiratory complex I, and localized within the thylakoid membranes of plants and cyanobacteria (*6, 7*). In land plants, the NDH comprises ∼30 subunits encoded in both the nucleus and the chloroplast (*8, 9*). It functions as a Fd-PQ oxidoreductase, transferring electrons from Fd to plastoquinone PQ during CEF, and as a protonmotive pump, translocating protons from the stroma into the lumen of the thylakoids (*2*). Although the activity of the NDH complex has been studied for decades, the physiological role of the complex in plants is still a broad topic for investigation and discussion. Tobacco plants lacking NDH have subtle phenotypes under standard growth conditions, while more pronounced mutant phenotypes can be seen under certain stress conditions (*10, 11*). The entire complex has been lost in several evolutionary lineages, some of which live in environments, or have energetic requirements, which one would expect to require maintenance of the complex [discussed in (*12*)]. Comparisons of NDH to its homologous respiratory complex may offer some ideas on the functions that the NDH complex may have in thylakoids. For example, the protonmotive nature of NDH allows for the thermodynamic reversibility of the complex, in that NDH can hypothetically oxidize PQH_2_ and transfer the electrons to Fd using *pmf* to drive the electrons energetically uphill [e.g., (*13, 14*)]. However, while this reverse reaction should be thermodynamically possible (*2*), it has yet to be demonstrated.

In order facilitate a structure-function investigation of NDH activity *in vivo*, we used chloroplast transformation in the model plant tobacco (*Nicotiana tabacum*) to modify a chloroplast genome-encoded proton channel-forming subunit of the NDH complex. Specifically, we mutated two conserved residues of the NdhF subunit known to be required for proton pumping (K240 and E152). We also attempted to delete the three C-terminal transmembrane helices that comprise the atypical helix spanning the membrane domain that is thought to act as a piston to drive proton translocation in complex I (*15*).

## Results

### Generation of Tobacco Plants with Mutated ndhF Genes

The subunits forming the proton channels of NDH are encoded entirely by the chloroplast genome. Therefore, modification of the corresponding genes requires chloroplast transformation. In this study, we focused our investigation on the NdhF subunit encoded within the small single copy region of the *Nicotiana tabacum* chloroplast genome (*16*). We mutated two residues that are conserved throughout complex I evolution and are required for proton pumping, lysine-240 (K240) and glutamate-152 (E152; *N. tabacum* numbering) (*17, 18*). We chose amino acid substitutions that are expected to abolish proton pumping, while maintaining similar side chain bulkiness: K240M and E152Q. Additionally, we generated a deletion mutant, in which the C-terminal transverse helix of NdhF associated with conformational change-induced proton pumping was removed by introducing a stop codon at amino acid position 645 (W645^*^).

For construction of our transformation vectors, we took advantage of the chloroplast genomic structure to facilitate post-transformation removal of the selectable marker cassette, a chimeric *aadA* gene conferring spectinomycin resistance. The chloroplast genome can take multiple configurations with regards to the inverted repeats (IR_A_ and IR_B_) and the relative orientations of the single copy regions (*19*). Due to flip-flop recombination of IR_A_ and IR_B_ (*16*), we expected to see four different conformations of the chloroplast genome after transformation. The two wild-type conformations (Figure 1A,B), and two conformations with the *aadA* cassette integrated into a BglII restriction site in the IRs (close to the border with the small single copy region). This integration occurs in both *ycf1* and *orf350*, a (non-functional) truncated fragment of *ycf1* residing in IR_B_ (Figure 1C,D). Due to the essentiality of *ycf1* (*20*), the transformed chloroplast genome will be maintained in a heteroplasmic state under spectinomycin selection, as configuration D (Figure 1D) would be lethal. Gene conversion between IR_A_ and IR_B_ (*21*) allowed us to screen for plants homoplasmic for the desired point mutations, and then to segregate out the *aadA* cassette from the chloroplast genome. To this end, seeds harvested from heteroplasmic transplastomic plants were germinated on selective medium containing 500 mg/l spectinomycin, and the white seedlings (sensitive to spectinomycin) were transferred to fresh media without selection and allowed to green.

**Figure 1.**
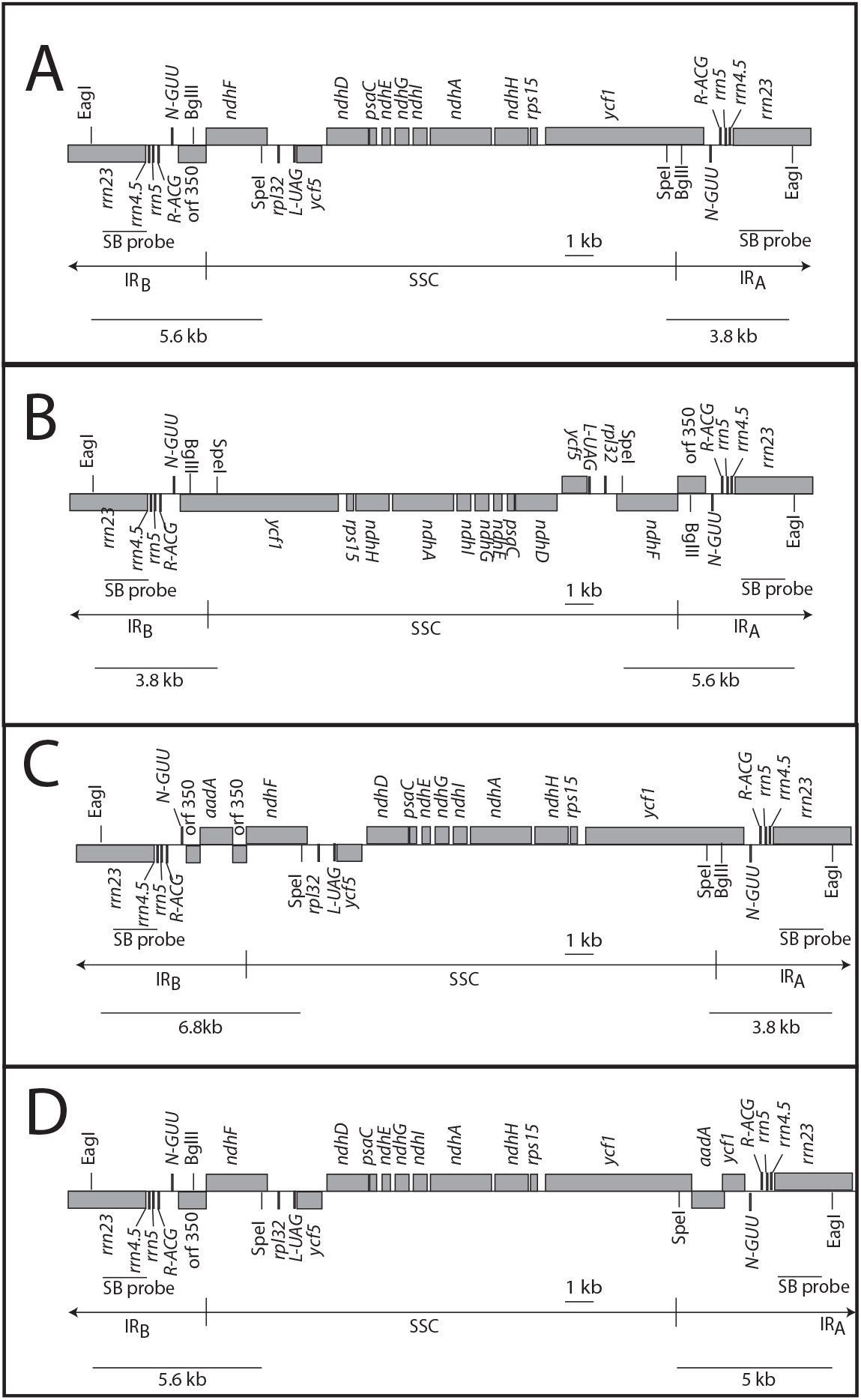
Genome conformations of chloroplast transformants generated in this study. (A) Wild type (WT) conformation 1 (*ycf1*-IR^A^). (B) WT conformation 2 (*ycf1*-IR^B^). (C) Transformed plastid genome conformation 1 (*aadA-orf350*-IR^B^). (D) Transformed plastid genome conformation 2 (*aadA-ycf1*-IR^A^). The radiolabeled probe used for Southern blot analysis is indicated (SB probe). IR^A^: inverted repeat A, IR^B^: inverted repeat B, SSC: small single-copy region of the plastid genome. Expected RFLP fragment sizes from a SpeI and EagI digest are indicated below each configuration.

Figure 1A-D illustrates the expected genome conformations in the transplastomic plants and the resulting restriction fragment sizes in RFLP (restriction fragment length polymorphism) analyses. Figure 2A shows a corresponding Southern blot analysis of green and white seedlings segregating from heteroplasmic parent plants. Wild-type plants and mutants in which the *aadA* cassette has been lost would give rise to hybridizing bands of 5.6 kb (*ndhF*-IR_B_) and 3.8 kb (*ycf1*-IR_A_), respectively (Figure 1A,B). Heteroplasmic mutants still containing the *aadA* cassette would produce the wild-type banding pattern and two additional bands corresponding to *ndhF*-*aadA*-IR_B_ and *ycf1*-*aadA*-IR_A_ at 6.8 kb and 5 kb, respectively (Figure 1C,D). In plants resistant to spectinomycin (green seedlings), we see bands for all four expected genome configurations (generated from ongoing flip-flop recombination of the inverted repeat), confirming the heteroplasmic state of the spectinomycin-resistant transplastomic lines. By contrast, the antibiotic-sensitive plants (white seedlings), show only the two bands characteristic of the *aadA*-free (wild type-like) genome. After greening on medium without antibiotic, these plants were grown to maturity and allowed to set seeds. DNA was extracted from a pool of seedlings (>20) and sequenced to assay for homoplasmy of the point mutations in *ndhF*. Figure 2B shows the sequencing chromatograms for the K240M mutations (AAA→ ATG) in lines 9 and 16, the E152Q mutations (GAA→ CAA) in lines 12 and 13, and the W645^*^ mutation (TGG→TAA) in lines 11 and 48. The absence of peaks for the wild-type nucleotide, together with the Southern blot results (Figure 2A), indicated homoplasmy for the point mutations and lack of the selectable marker gene cassette. For each point mutation, two independently generated homoplasmic marker-free transplastomic lines were chosen for in-depth characterization.

**Figure 2.**
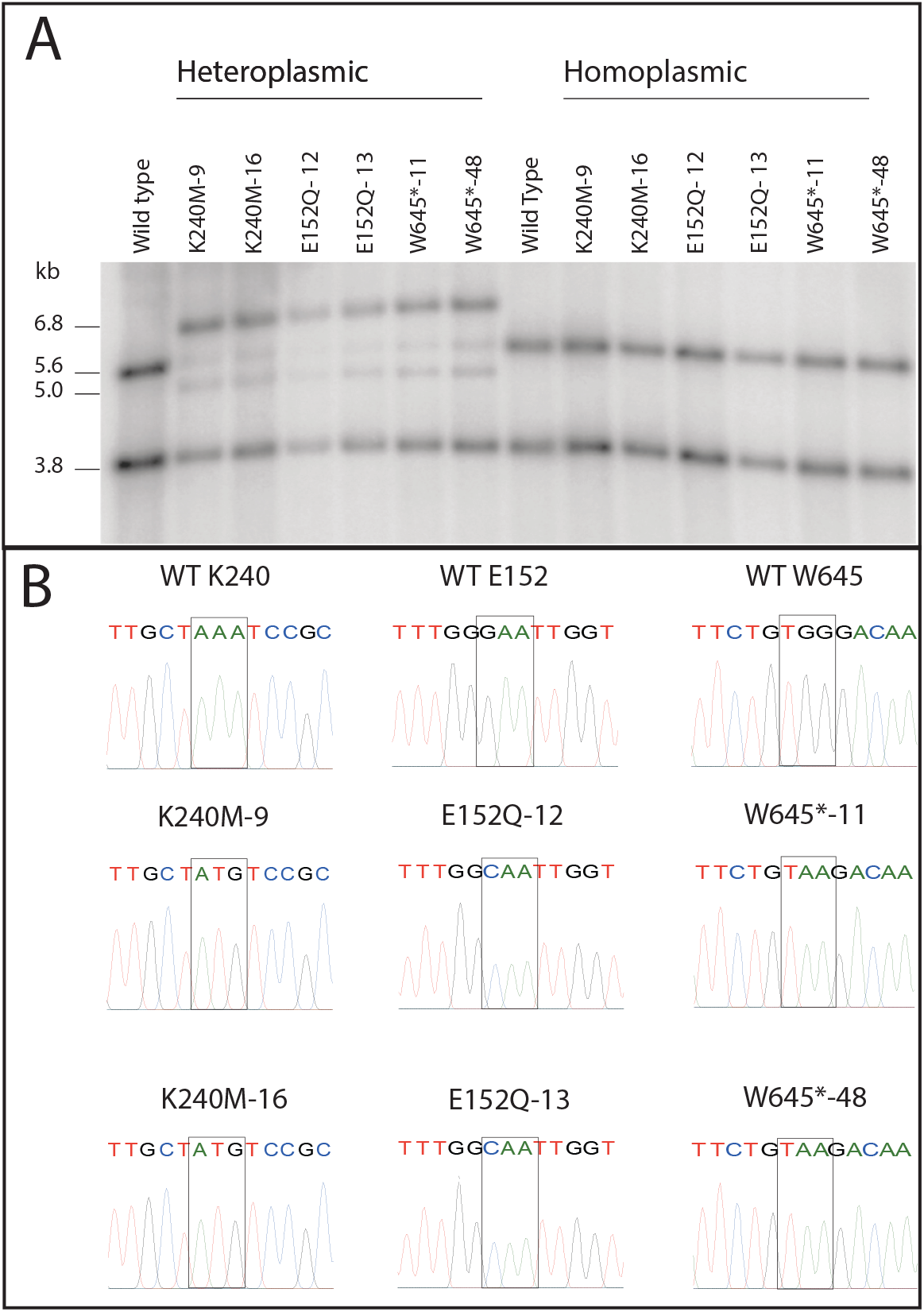
Identification of homoplasmic transplastomic tobacco lines with point mutations in the *ndhF* gene. (A) Southern blot analysis of heteroplasmic and homoplasmic mutant lines obtained from the segregating progeny of a heteroplasmic parent plant. The heteroplasmic plants harbor all 4 genome conformations illustrated in Figure 1 (A-D), the homoplasmic lines have lost the selectable marker gene, while retaining the (homoplasmic) point mutations in the *ndhF* gene. (B) Conformation of the homoplasmic state of the point mutations in *ndhF* by DNA sequencing. DNA was extracted from pooled seedlings, and the PCR-amplified *ndhF* gene was sequenced. Boxes indicate the mutated codons.

### NDH Complex Content and Assembly

To identify possible changes in NDH protein complex accumulation, we investigated the NDH complex abundance in thylakoid membranes of the transplastomic mutants. To this end, the accumulation of two diagnostic subunits of the complex was determined with specific antibodies: NdhB and NdhH (Figure 3). As a control for equal quality of the thylakoid preparations and equal loading, immunoblots were probed with antibodies against Lhcb2, a subunit of the ligh-harvesting antenna of photosystem II. In addition, a previously generated NDH knockout line that had been obtained by inactivation of the *ndhF* gene [*ΔndhF* (*22*)] was included as a negative control. The immunoblot analyses revealed lower contents of the NDH subunits NdhB and NdhH in the K240M and E152Q mutant lines compared to the wild type. The K240M and E152Q substitution lines accumulated 25 and 50% of wild-type NDH content, respectively. The point mutations led to a decrease in NdhB and NdhH subunit content, which appears to be stoichiometric, suggesting that the complex is still assembled but with lower efficiency. Similar to the *ndhF* knockout mutant, the W645^*^ lines lacked accumulation of the NdhB and NdhH subunits to detectable levels.

**Figure 3.**
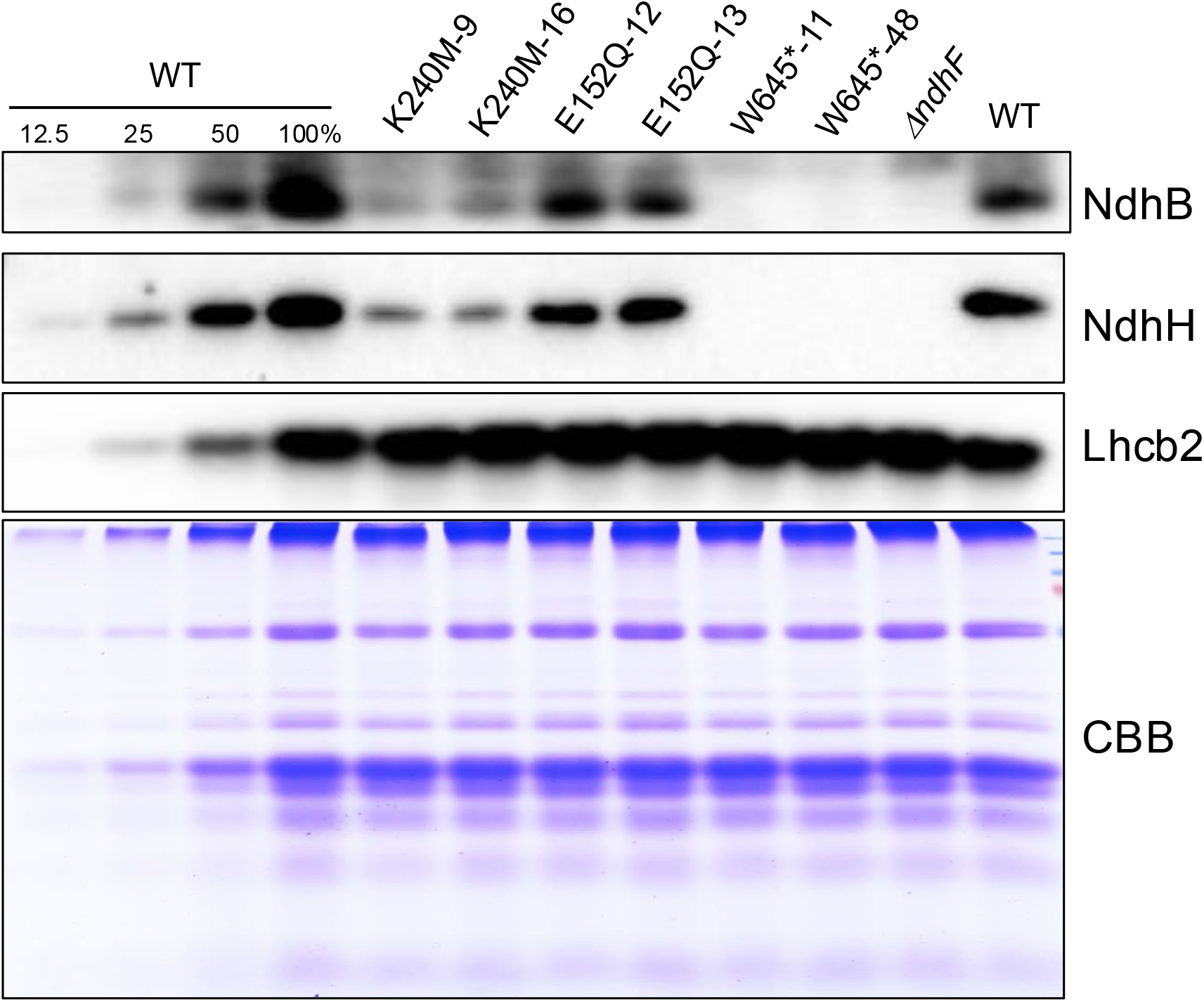
Accumulation of NDH subunits in thylakoid samples from the wild type (WT), the ndhF point mutation lines (K240M-9, K240M-16, E152Q-12, and E152Q-13), the truncated lines (W645*-13 and W645*-48), and the *ndhF* knockout mutant (*ΔndhF*). Thylakoid samples equivalent to 6 μg of chlorophyll (100%) were resolved by electrophoresis in a 12.5% SDS-PAA gel. The immunodetection was conducted with the antibodies anti-NdhB and anti-NdhH. As controls for equal loading, a blot was probed with an anti-Lhcb2 antibody, and the Coomassie brilliant blue (CBB) staining of the protein gel is shown below the blots.

To determine if the remaining NDH complex in our mutants was properly assembled into PSI-NDH supercomplexes, we conducted blue-native polyacrylamide gel electrophoresis (BN-PAGE, Figure 4A) experiments followed by immune detection of NdhH (Figure 4B). In wild type plants the NdhH subunit is associated with three different complexes. These correspond to the NDH complex not associated to PSI (around 700 kDa and migrating slightly below PSI), a second complex not yet characterized (indicated with the red asterisk in Figure 4B), and the PSI-NDH supercomplex (Figure 4A). In the proton channel point mutants K240M and E152Q, formation of the PSI-NDH supercomplex is still observed, but the accumulation of the free NDH complex is reduced in these lines. Consistent with our immunoblots, no accumulation of NDH complex can be detected in the W645^*^ mutant lines and in the *ndhF* knock-out mutant (*ΔndhF*). These results suggest that the K240M and E152Q substitutions do not interfere with NDH assembly with PSI. In addition, since the deletion of the transverse helix of NdhF in the W645^*^ mutants leads to complete loss of the PSI-NDH supercomplex in the first dimension, with no evidence for NdhH accumulation in the free NDH complex or in the PSI-NDH supercomplex, it may be that the presence of the transverse helix in NdhF is essential for complex assembly and/or stability.

**Figure 4.**
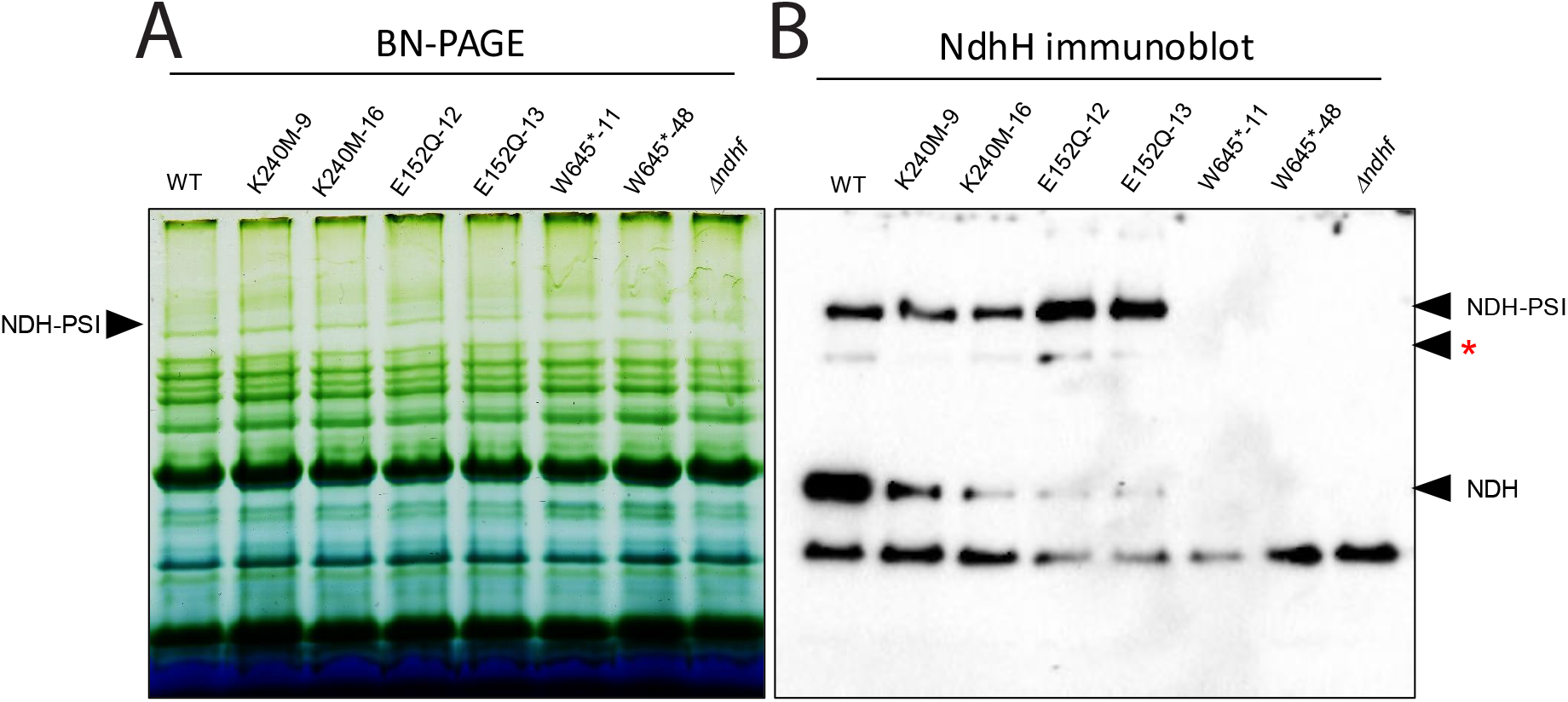
Detection of NDH supercomplexes in thylakoid samples from the wild type (WT), the *ndhF* point mutation lines (K240M-9, K240M-16, E152Q-12, and E152Q-13), the truncated lines (W645*-11 and W645*-48), and the *ndhF* knockout line (*ΔndhF*). (A) First-dimension separation of thylakoidal protein complexes using blue-native polyacrylamide gel electrophoresis (BN-PAGE). (B) Immunodetection of the NDH-containing complexes and supercomplexes using the anti-NdhH antibody. The red asterisks marks the accumulation of an uncharacterized complex containing NdhH.

### NDH Activity in Transplastomic ndhF Mutants

In order to determine if the NDH complex still retained plastoquinone reductase activity in our NDH mutants, we used a modified approach to visualize the post-illumination fluorescence rise associated with NDH reduction of the PQ pool (discussed in (*2*)). Leaves were illuminated for 5 minutes in 115 μmol photons m^-2^ s^-1^ and the fluorescence was normalized to the maximum fluorescence yield during a 500 ms saturating flash after 10 minutes of dark adaptation. A saturation flash was applied every 60 s during the illumination phase.

Figure 5A shows representative data from the full F_M_ normalized fluorescence trace of the wild type. Figure 5B shows a close-up of the normalized fluorescence after the light to dark transition. In the wild type, after switching the actinic light off, the fluorescence yield dropped and then rose again in a typical manner associated with PQH_2_ oxidation and then PQ reduction by NDH. In the *ndhA* knockout mutant [*ΔndhA*, (*23*)], and the *ndhF* mutants generated in this study, the fluorescence signal during the dark interval after illumination did not rise. After 50 s in the dark, a 50 second far-red (FR) pulse was applied to re-oxidize the PQ pool, followed by another 150 s dark interval. This FR pulse followed by dark was repeated 2 additional times. In the wild type, fluorescence increased during each dark interval following a FR pulse, with a decrease in slope occurred after each subsequent FR pulse. This behavior may be due to depletion of stromal reducing equivalents during the duration of the experiment. In the mutants there is an increase in fluorescence during the FR pulse, which may be an artifact due to excitation of PSII by the FR source, which quickly drops after FR illumination is terminated and the PQ pool oxidizes. The slope of the fluorescence rise after each FR pulse in the mutants is either severely impaired or abolished. It is likely that any small amount of post-illumination rise seen in the mutant lines is due to a slight actinic effect of the measuring beam. Taken together, these data show that all our *ndhF* mutants have lost the PQ reductase activity of the NDH complex, regardless of protein or supercomplex content.

**Figure 5.**
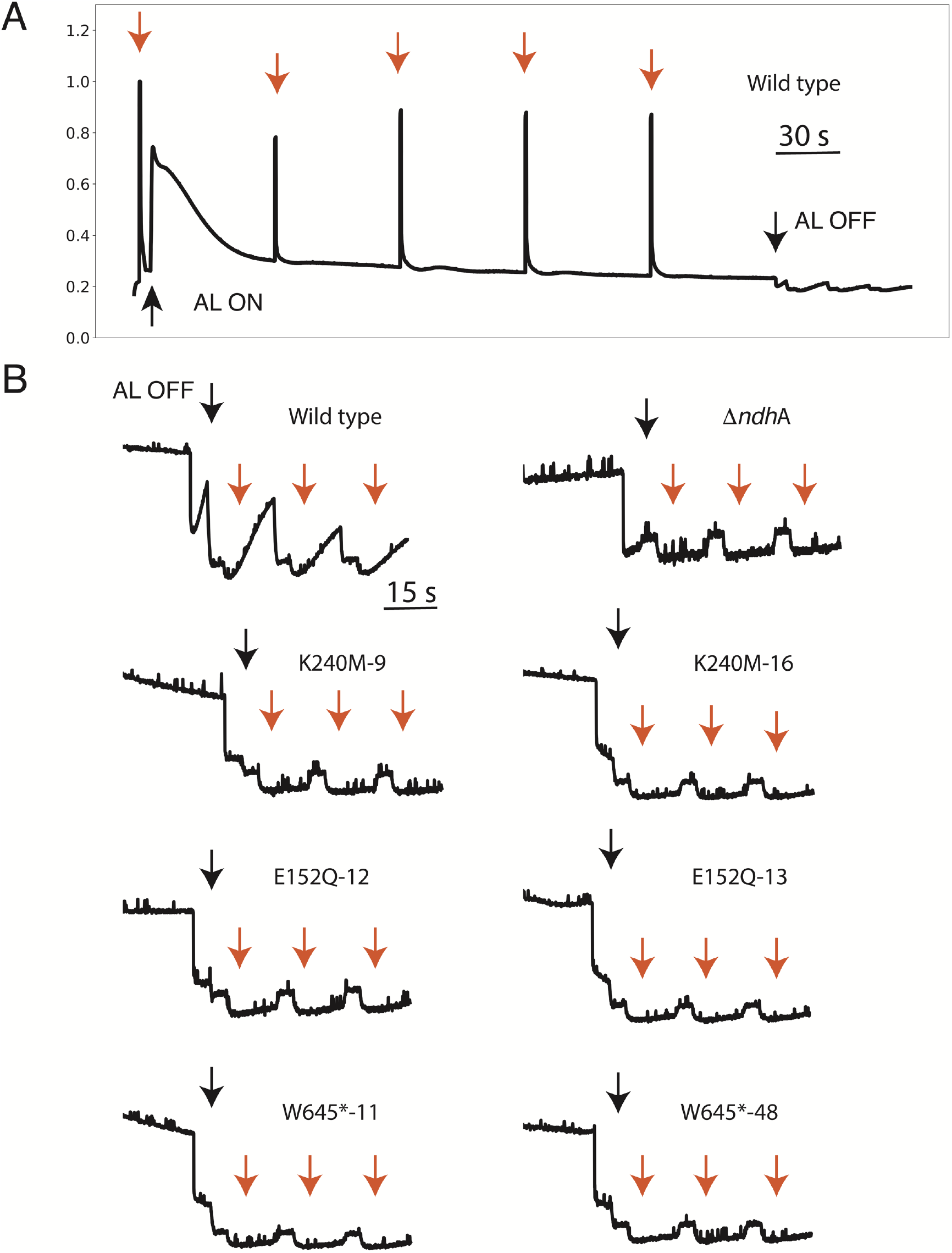
Post-illumination fluorescence rise kinetics. A) representative wild type trace. Red arrows indicate application of a 500 ms saturation flash, black arrows indicate appilication and cessation of actinic light (AL). B) Close-up of post-illumination fluorescence phase. Black arrows indicate cessation of actinic beam, red arrows indivate application of a 30 s far-red pulse. Mutant lines are indicated above each representative trace, with the ndhA knockout line (ΔndhA) serving as a negative control. Data were normalized to the dark-adapted fluorescence maximum during a saturation flash.

### Steady-State Photosynthetic Phenotypes of Transplastomic NDH Complex Mutants

The post-illumination fluorescence rise relies on priming the system to favor dark reduction of the plastoquinone pool. While this has been immensely effective in determining relative NDH activity [e.g., (*24*)], it is difficult to quantify and may not be directly applicable to the state of the system in the light. Therefore, to determine the impact of our mutations on steady-state photosynthesis, we used chlorophyll *a* fluorescence and the electrochromic shift (ECS) of the carotenoid absorbance spectrum around 520 nm. Chlorophyll fluorescence allows for non-invasive photosynthetic measurements related to electron transfer and photoprotection, while ECS is reflective of the membrane potential, and is useful as a probe of the thylakoid proton circuit.

Due to the protonmotive nature of the NDH complex, we might expect defects in this route of CEF to impact both electron transfer and downregulation of light harvesting at PSII, processes that are sensitive to feedback regulation via pH decreases in the thylakoid lumen. Electron transfer originating at PSII (linear electron transport, LEF) is modulated by ΔpH-induced *q*_E_ and PCON. We might expect that losing a proton-coupled electron transfer route in the thylakoid (NDH-CEF) would lead to an increase in LEF due to a relaxation of feedback regulation. Instead, in the NdhF-K240M, E152Q, and W645^*^ mutants, LEF values were similar to the wild type (Figure 6A, C, and E). Total NPQ was also similar or slightly elevated at the highest light intensity in the NDH mutants (Figure 6B, D, F). Additionally, total *pmf*, as estimated by ECS_*t*_, was similar or slightly decreased (Figure 7A, C, E), indicating there was little impact on the overall amplitude of *pmf*. Interestingly, trans-thylakoid proton conductivity (i.e., the rate constant of proton efflux, *g*_H_^+^) is increased in the NDH mutants (Figure 7B, D, F). This increase in the NDH mutants of *g*_H_^+^ is most noticeable at lower light intensities and may be the cause for the decreased total *pmf* seen in the NDH mutants.

**Figure 6.**
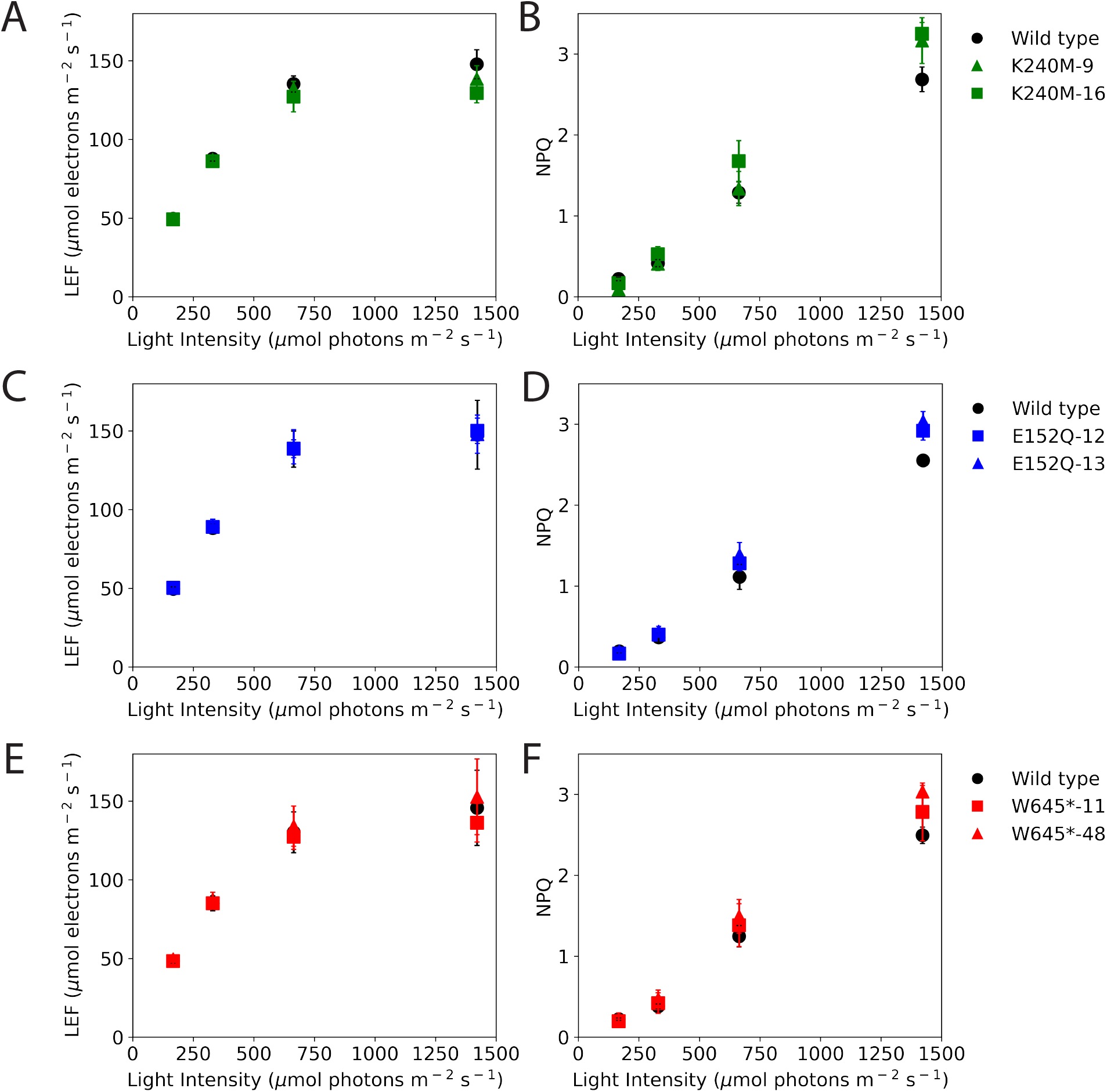
Light-response curves of chlorophyll a fluorescence parameters in *ndhF* mutant lines. Linear electron flow (LEF; A, C, D) and non-photochemical quenching (NPQ; B, D, F) are shown for the wild type (black circles), the *ndhF* point mutation lines K240M (A and B) -9 (green triangles) and -16 (green squares), E152Q (C and D) -12 (blue squares) and -13 (blue triangles), and the truncated NdhF lines W645* (E and F) -11 (red squares) and -48 (red triangles).

**Figure 7.**
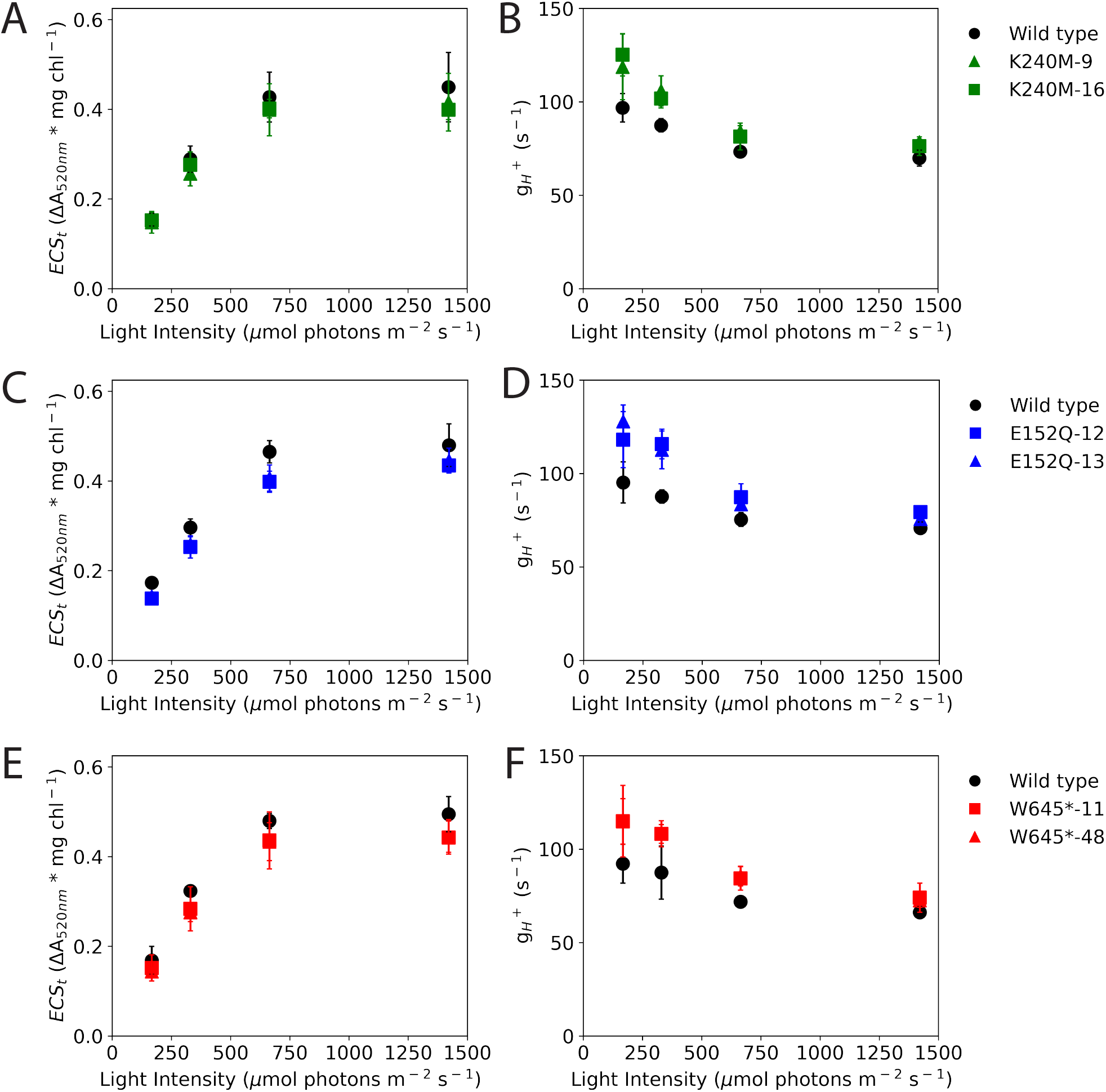
Light-response curves of electrochromic shift-derived parameters in *ndhF* mutant lines. Total light-driven protonmotive force as estimated by the total electrochromic shift at 520 nm from light to dark (ECS_*t*_; A, C, D) and the trans-thylakoid proton conductivity (*g*_H_ ^+^; B, D, F) are shown for the wild type (black circles), the *ndhF* point mutation lines K240M (A and B) -9 (green triangles) and -16 (green squares), E152Q (C and D) -12 (blue squares) and -13 (blue triangles), and the truncated NdhF lines W645* (E and F) -11 (red squares) and -48 (red triangles).

The post-illumination fluorescence measurements indicated that NDH activity was likely lost, regardless of protein complex content. Next, we wanted to quantify the loss of proton translocation into the lumen. Since LEF has a fixed ratio of 3H^+^/e^-^ and NDH-mediated CEF has a ratio of ∼4H^+^/e^-^ [discussed in (*2, 3*)], if a substantial contribution of NDH mediated CEF was lost, we would see a decrease in the ratio of trans-thylakoid proton efflux (*v*_H_^+^) to LEF (Figure 8A, C, E). As described for Arabidopsis NDH loss-of-function mutants before (*25*), there is no apparent decrease in the slope of this function in any of our tobacco *ndhF* mutants outside of the level of experimental noise. We also took a secondary method for measuring CEF by comparing the ratio of *pmf* (ECS_*t*_) to *pmf* generated by LEF alone (*pmf*_LEF_) (*26*) (Figure 8B, D, F). If CEF was decreased, we would expect to see a decrease in the slope of this relationship. Again, we see only subtle decreases within the expected noise level in the dependency of ECS_*t*_ and *pmf*_LEF_ for any of our mutations regardless of protein content. These data are in agreement with data from the characterization of Arabidopsis mutants that are completely deficient in NDH complex activity (*25*).

**Figure 8.**
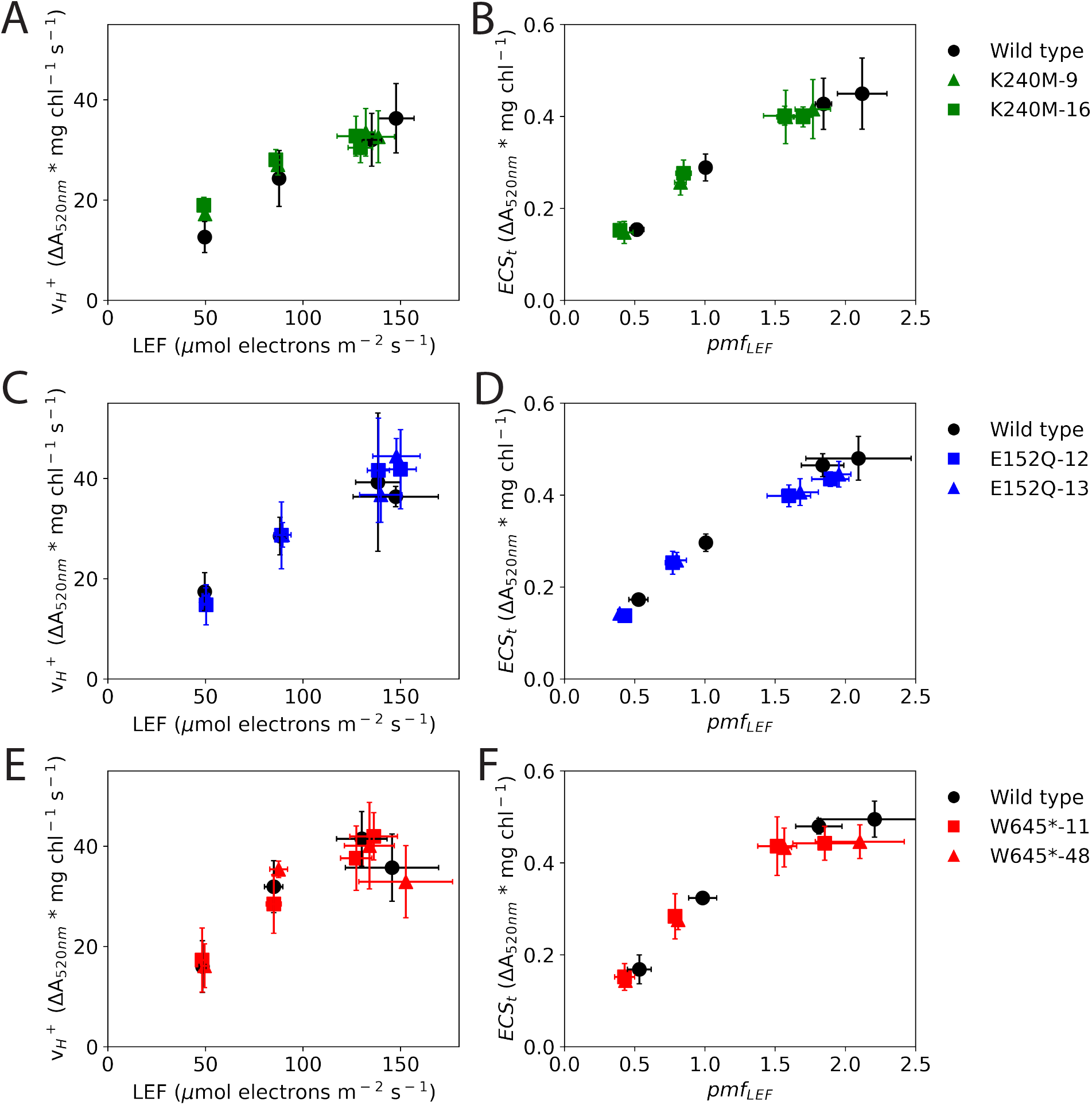
Estimation of changes in cyclic electron flow rates in the *ndhF* mutant lines. Trans-thylakoid proton flux (*v*_H_ ^+^) as a function of linear electron flux (LEF) (A, C, D), and total light-driven protonmotive force (*pmf*) as estimated by the total electrochromic shift at 520 nm from light to dark (ECS_*t*_) as a function of *pmf* generated by LEF alone (*pmf*_LEF_) (B, D, F) are shown for the wild type (black circles), the *ndhF* point mutation lines K240M (A and B) -9 (green triangles) and -16 (green squares), E152Q (C and D) -12 (blue squares) and -13 (blue triangles), and the truncated NdhF lines W645* (E and F) -11 (red squares) and -48 (red triangles).

NDH mutants have in the past been described as having increased NPQ [e.g., (*27*)]. As *pmf* is comprised of both a ΔpH and a Δψ component, of which the ΔpH component is regulatory, all else being equal, we would expect that the slight decreases in ECS_*t*_ we see in our mutants would lead to a slight decrease in the ΔpH-dependent component of NPQ, *q*_E_. However, we see an increase in NPQ (Figure 6B, D, F), of which *q*_E_ is the largest component, suggesting an altered sensitivity of *q*_E_ to *pmf*. To demonstrate this effect, we plotted *q*_E_ as a function of ECS_*t*_ (Figure 9). All of our NDH mutations showed an increase in the sensitivity of *q*_E_ to *pmf*.

**Figure 9.**
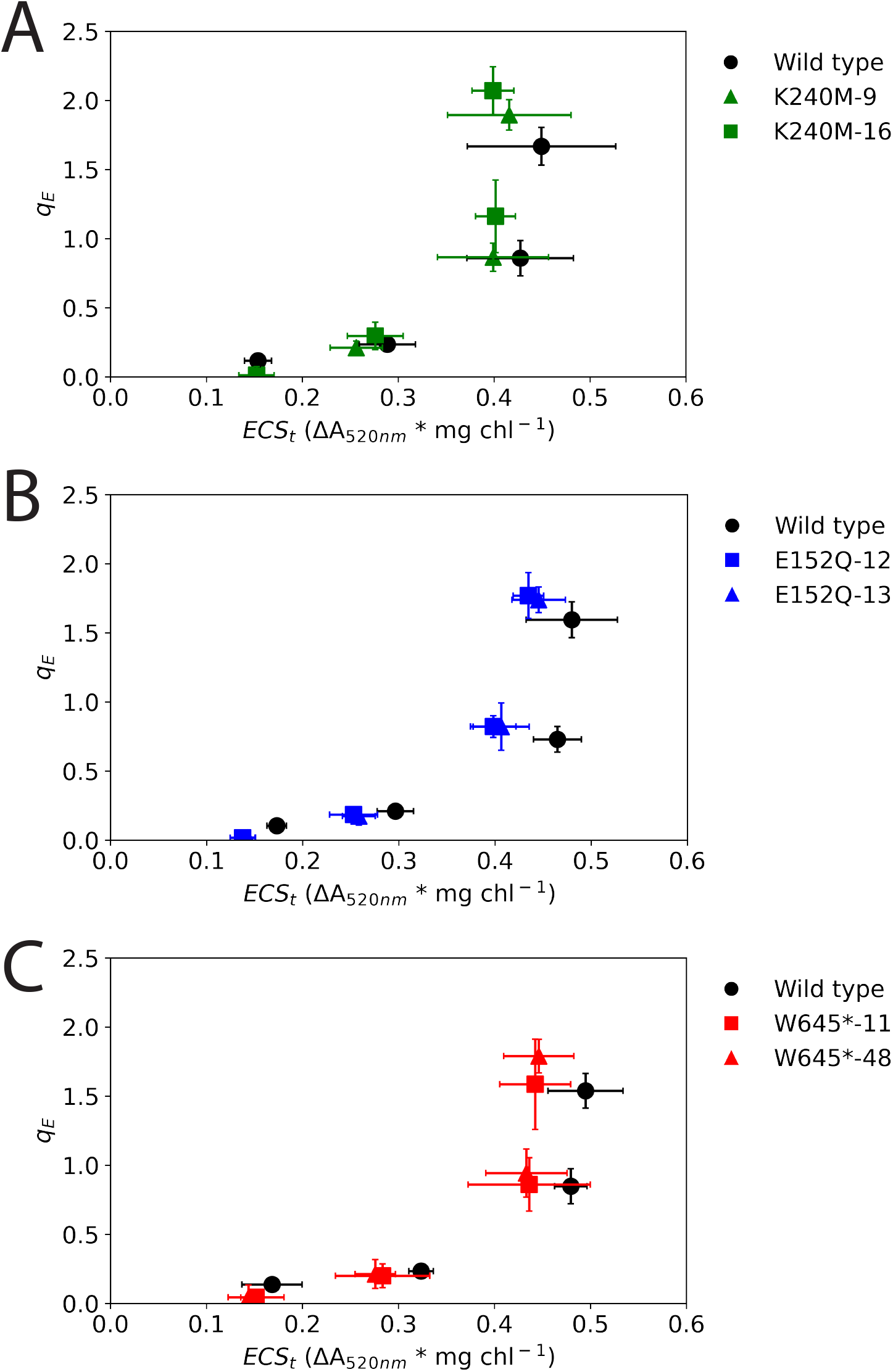
Sensitivity of pH-dependent antenna downregulation to protonmotive force (*pmf*). Energy-dependent quenching (*q*_*E*_) as a function of light-driven *pmf* as estimated by the total electrochromic shift at 520 nm from light to dark (*q*_*E*_ is shown for the wild type (black circles), the *ndhF* point mutation lines K240M (A) -9 (green triangles) and -16 (green squares), E152Q (B) -12 (blue squares) and -13 (blue triangles), and the truncated NdhF lines W645* (C) -11 (red squares) and -48 (red triangles)

### Dependency of PSI-NDH supercomplex formation on NDH accumulation

It has been proposed that the NDH complex requires interaction with PSI for both stability and function (*28, 29*). Since the W645* mutants lost detectable NDH protein content, we hypothesized that this could be explained by loss of interaction with PSI due to deletion of the atypical transverse helix, which may be an important interaction point of the PSI-NDH supercomplex (*30*). To test this hypothesis, we looked at NDH complex accumulation in a transplastomic knockout mutant of *ycf3*, a gene that is required for the assembly of PSI (*31–33*). Additionally, to test whether complex accumulation is impacted by loss of any component of the electron transport chain, we analyzed NDH complexaccumulation in transplastomic knockout mutants of *petN*, an essential subunit of the bf complex (*34*) *psbD*, a core subunit (D2) of PSII (*35, 36*), and *ndhCKJ*, an operon comprising three genes for subunits of the NDH complex (*24*). As photosynthesis is not essential under heterotrophic conditions, plants can be grown without a complete electron transport chain when supplemented with sucrose. Figure 10A shows immunoblot analyses of these mutants with antibodies against AtpB, NdhH, and PSAD. Despite loss of PSAD in the *ycf3* knockout mutant, the NdhH protein still accumulates to wild type-like levels, thus invalidating the hypothesis that PSI-NDH complex formation is required for NDH accumulation. Figure 10B shows immunoblots against PetA and PsbD. With the exception of the *petN* mutant, there is accumulation in our photosynthetic mutants of all complexes probed relative to AtpB. While the *petN* mutant has no defect in photosynthetic complex accumulation other than the *bf* complex in very young leaves grown under extreme low-light conditions (*34*), it may be possible that the PsbD subunit of PSII is secondarily reduced due to photodamage accumulation occurring in *petN* mutant plants under our growth conditions.

**Figure 10.**
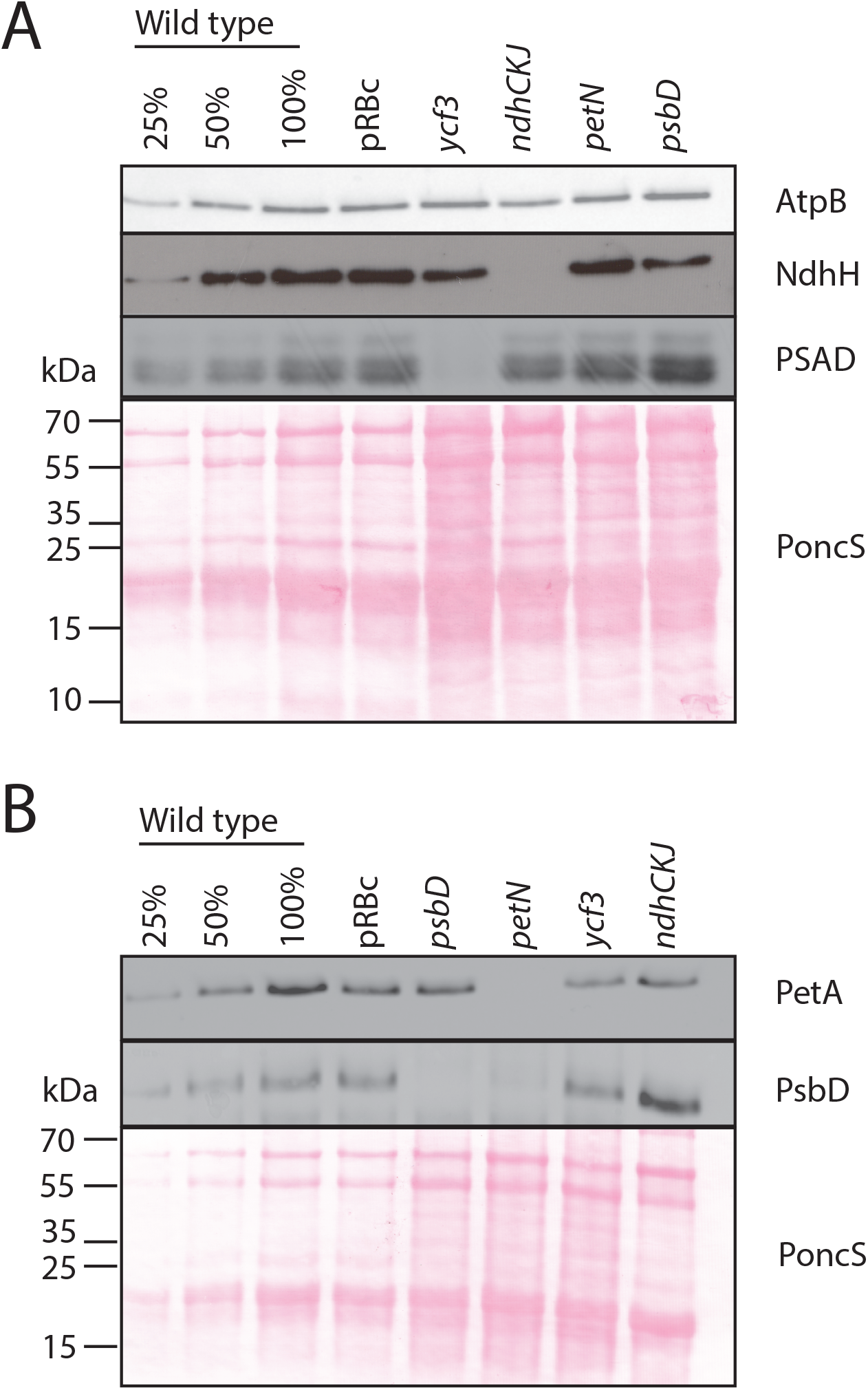
Accumulation of representative subunits of the main components of the electron transport chain in knockout lines of the main complexes of the electron transport chain. (A) immunodetection of AtpB, NdhH, and PSAD (B) immunodetection of PetA and PsbD. PRB8 #E15 (pRBc) was used as a control plant for the presence of the *aadA* cassette. Due to differences in chlorophyll:protein content, lanes were loaded to achieve equal accumulation of AtpB.

## Discussion

The NDH complex is enigmatic in its function within the thylakoid membrane. Despite its relatively high level of conservation, mutants lacking the complex in model plant species have no growth defects unless grown under stressful conditions [e.g., (*10*)]. Additionally, photosynthetic phenotypes in unstressed plants are subtle, and are not explained easily by a loss of NDH-mediated CEF [e.g., (*10, 27*)]. As the chloroplast NDH complex is homologous to respiratory complex I, we used what is known from complex I to try and explain these subtle phenotypes for a more complete understanding of the function of NDH within the context of photosynthesis. In this study, we focused on NdhF, one of the proton channel-forming subunits encoded by the chloroplast genome. By modifying protonmotive residues, we wanted to alter the H^+^/e^-^ transfer ratio of the complex. However, mutation of conserved residues within NdhF (K240 and E152), and deletion of the transverse helix predicted to participate in proton shuttling led to loss of plastoquinone reductase activity (Figure 5), indicating that proton translocation and electron transfer are tightly coupled in the chloroplast NDH complex. This is in stark contrast to bacterial systems where truncating or eliminating the entire NdhF homolog, NuoL, fails to abolish electron transfer (*37*).

Since our mutant plants still retained varying amounts of the NDH complex, we were able to investigate the impact on the electron transport chain when complex activity is lost, but the interaction of the complex with PSI is retained (Figure 4). While all of our NDH mutations led to loss of the plastoquinone reductase activity (as evidenced by chlorophyll fluorescence measurements in the post-illumination phase; Figure 5), in the steady state, there is no persuasive evidence for the loss of the forward Fd:plastoquinone reductase activity [Figure 8 and e.g., (*25*)], and no evidence for the loss of the reverse reaction. However, increased *g*_H+_ conductivity in these mutants could indicate an ATP deficit, thus indirectly suggesting a loss in the forward reaction of the NDH complex. Thus, the forward reaction could augment ATP production at levels that are outside the sensitivity of our methodology.

In addition to the increased *g*_H+_, our collection of NdhF mutants also show increased sensitivity of *q*_E_ to *pmf*. This increased sensitivity of *q*_E_ may arise from two functions associated with the NDH complex. First, the loss of an alternative electron transfer pathway may lead to an altered chloroplast redox state, which in turn may impact redox cofactors involved in the regulation of the xanthophyll cycle (*38, 39*). However, if the Fd:PQ reduction route of electron transfer through the NDH complex were completely lost, one might expect a more reduced stroma, leading to inhibition of violaxanthin de-epoxidase and the loss of *q*_E_ sensitivity to ΔpH. Second, a change in distribution of *pmf* into an increased fraction of ΔpH and a decreased fraction of Δψ would lead to an increased sensitivity of *q*_E_ to total *pmf*. This change in *pmf* partitioning could occur due to the antiporter legacy of the complex [i.e., its complex I ancestry; (*40, 41*)]. Evolutionarily, complex I arose from association of several H^+^/Na^+^ antiporter proteins. In several lineages of complex I evolution, both prokaryotic and eukaryotic, the antiporter function has been retained and is associated with the NdhF homologs in these species (*41, 42*). If antiporter activity was maintained in the NDH lineage of complex I evolution, our NDH complex mutations in the NdhF subunit would be expected to abolish it, as NdhF is the subunit that is responsible for this activity. Defects in *pmf* partitioning that might arise due to the loss of antiporter activity could explain why an increase in the sensitivity of *q*_E_ to ΔpH is observed in our mutant lines without substantial changes in proton and electron fluxes.

In summary, in this study, we have shown that the proton and electron transfer reactions within the NDH complex are tightly coupled and that the photosynthetic phenotypes associated with the loss of NDH cannot be entirely explained by, and in some instances are conflicting with, loss of NDH mediated CEF. Future work should be directed towards elucidating a possible secondary function of the NDH complex not associated with meeting chloroplast ATP demands, such as the possible function of NdhF in luminal ionic regulation.

## Materials and Methods

### Plant growth and materials

All mutants used and generated in this work were in the *Nicotiana tabacum* cv. Petit Havana background. Leaf tissue for DNA, protein, and spectroscopic analysis was from plants grown as described in (*43*) with the exception of tissue used in Figure 10. For Figure 10, plants were grown in tissue culture as described in (*34*).

### Construction of plastid transformation vectors

The region of the tobacco chloroplast genome containing the *ndhF* region was cloned as a 4347 bp fragment into plasmid pMCS5 (MoBiTec) between the PacI and PmeI restriction sites using a 3 fragment Gibson assembly protocol (*44*) with the primer pair P1 and P2 and the primer pair P3 and P4, generating plasmid pDDS008 (Supplemental Table 1). To conduct the site-directed mutagenesis, a 748 bp fragment of *ndhF* was amplified by PCR using primer pair P5 and P6, digested with NdeI and SpeI, and cloned into pMCS5 generating pDDS002. For generation of the K240M mutation by site-directed mutagenesis, the plasmid pDDS002 was amplified first with primer pair P7 and P8, introducing an AAA → AAG mutation with the primer sequence, and then amplified again with primer pair P9 and P10 to generate the AAG→ ATG mutation (pDDS004). For the E152Q mutation, pDDS002 was amplified with primer pair P11 and P12 to generate the GAA → CAA mutation (pDDS006). After each amplification, the product was treated with DpnI to remove the plasmid template, and subsequently transformed into *E. coli* for replication and confirmation by DNA sequencing. To introduce the mutated sequences into vector pDDS0008, ndhF fragments were amplified from pDDS004 and pDDS006 via PCR using primers P5 and P6, digested with NdeI and SpeI, and ligated into the similarly cut pDDS008 between the NdeI and SpeI sites generating plasmids pDDS010 and pDDS011, respectively.

To generate a stop codon at position W645, the chloroplast DNA region to be included in the transformation vector was assembled into pMC5 (digested with PacI and PmeI) using a 3 fragment Gibson assembly protocol and primers harboring the TGG—>TAA mutation, generating plasmid pDDS009. The primer pairs were P13/P14 and P15/P16 (Supplementary Table 1). Finally, the selectable marker gene cassette was amplified from pDK308 (*45*) using the primer pair P17 and P18. The cassette contains the *aadA* gene conferring spectinomycin resistance flanked with the psaA promoter and *atpB* terminator from *Chlamydomonas Reinhardtii*, and was inserted into all three plasmids (pDDS010, pDDS011, and pDDS009) by Gibson assembly at the BglII restriction site. The resulting vectors (pDDS0013, pDDS0014, and pDDS0016, respectively) were used for biolistic transformation.

### Chloroplast transformation

*Nicotiana tabacum* (cv. Petit Havana) leaves were biolistically transformed as described previously (*46, 47*). Spectinomycin-resistant lines were selected on regeneration medium containing 500 mg/L spectinomycin. Resistant shoots were analyzed for presence of the point mutations by PCR amplification and restriction digest. For E152Q, the GAA → CAA mutation introduced a restriction site for MfeI. The region was amplified using primer pair P19 and P20 (Supplementary Table 1), and digestion gave a 785 bp band for the wild-type sequence, and two bands of 432 bp and 353 bp for the mutated sequence. For the K240M and W645^*^ lines, dCAPS markers with polyA extensions were used to introduce a restriction site for identification of chloroplast transformants that harbor the desired mutations. For K240M, primer pair P21 and P22 introduced an MseI restriction site in the wild-type sequence, resulting in 2 bands of 34 bp and 285 bp for the mutated sequence, and 3 bands of 34 bp, 57 bp and 255 bp for the wild-type sequence. For W645^*^, primer pair P23 and P24 introduced a HaeIII restriction site into the wild-type sequence giving a 287 bp band and two bands of 230 bp and 57 bp for the mutant and wild-type sequences, respectively. All fragment differences were visible upon electrophoretic separation in a 2% agarose gel.

Tissue from shoots that were positive for the desired mutations (either appearing hetero- or homoplasmic) went through an additional round of regeneration and PCR-based screening for the desired mutations. Plants appearing to be homoplasmic for the mutation at this stage were placed into the greenhouse and allowed to set seeds. For removal of the selectable marker gene cassette, seeds from individual lines were sown on selection medium, white (antibiotic-sensitive) seedlings were removed from selection and allowed to regreen. Tissue samples from the growing seedlings were then analyzed by Southern blot and DNA sequencing for absence of the marker gene cassette and homoplasmic presence of the point mutations.

### Southern blot analysis

Total genomic DNA was isolated using the protocol described in (*48*), with the following modifications: Following the recovery of the aqueous phase after the chloroform-isoamyl alcohol extraction, samples were treated with RNase A for 30 minutes at 16°C and underwent a second chloroform-isoamyl alcohol extraction. The final pellet was resuspended in sterile deionized water. Total DNA (3 μg) was digested with the restriction enzymes EagI and SpeI overnight and run on a 0.7% agarose gel. The gel was then blotted by capillary action onto a positively charged nylon membrane (Hybond XL, Amersham Biosciences) overnight. The membrane was probed with a 655 bp fragment of the plastid 23S rRNA gene within the inverted repeat generated by PCR using the primer pair P25 and P26 (Supplementary Table 1), and labeled with [α-^32^P]dCTP by random priming (Multiprime DNA labeling kit; GE Healthcare).

### SDS-PAGE, BN-PAGE and western blot analysis

Crude chloroplasts were isolated as described in (*49*), with the addition of 10 mM ascorbate. The chloroplasts were then osmotically ruptured on ice for 10 minutes in > 5x excess buffer (10 mM HEPES, 5 mM MgCl_2_, 2.5 mM EDTA, 10 mM ascorbate, pH 7.6). Thylakoids were pelleted at 3000 x g and resuspended in the osmotic shock buffer and chlorophyll content was determined according to (*50*). Thylakoids were solubilized in Laemmli buffer supplemented with 20 mM DTT, and separated in a 12.5% SDS polyacrylamide gel (*51*). Proteins were then transferred to a PVDF membrane and probed with the antibodies anti-NdhB (Agrisera; cat. AS16 4064), anti-NdhH (Agrisera cat. AS16 4065) or LHCB2 (Agrisera cat. AS01 003) using the dilution recommended by the manufacturer.

Blue-native PAGE was conducted according to (*51*). Thylakoid samples were solubilized in 1% DDM and separated in a 6-12% native polyacrylamide gel. Then, the gel was incubated for 1 hour in Laemmli buffer containing 100 mM DTT and 8 M Urea. Finally, the proteins were transferred to a PVDF membrane and immunodecorated with the antibodies described above.

### Photosynthesis measurements

Plants were dark adapted for 30 minutes prior to experimentation. Saturation pulse chlorophyll *a* fluorescence yield parameters (F_o_, F_m_, F_s_, F_m_’’) were measured at actinic intensities from 100 to 1500 μmol photons m^-2^ s^-1^ as described in (*52*). Fm’ was calculated using a multiphase flash described in (*53*). These parameters were used to calculate photosynthetic efficiency (ϕ_II_), total non-photochemical quenching (NPQ), and exciton quenching (q_E_) as described in (*54*). Linear electron flow was calculated as in eq 1.

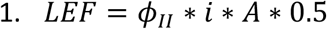

Where *i* is the actinic intensity, A is the absorptivity of the leaf [measured as described in (*55*)], and 0.5 is the assumed fraction of light absorbed by photosystem II.

The post-illumination chlorophyll fluorescence transient was measured in intact leaves as described in (*56*).

Electrochromic shift measurements were performed in the steady state by monitoring absorbance changes in a measuring pulse supplied with a green LED (Luxeon Rebel) filtered with a 520 nm bandpass filter (Intor). Amplitude-dependent parameters (*v*_H_^+^ and ECS_t_) were normalized to chlorophyll content [measured according to (*50*)]. ECS_*t*_ was taken as the total extent of the shift during a 300 ms dark interval, while *v*^H+^ was taken from the initial slope of the decay. The relative extent of *pmf* attributable to LEF (*pmf*_LEF_) was calculated as in eq. 2 (*26*).

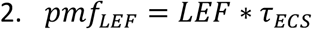

Where *τ*_*ECS*_ was calculated from a first order exponential fit of the decay during the light-to-dark interval.

## Abbreviations

ATP: adenosine triphosphate
*bf*: cytochrome *b*_6_*f* complex
BN-PAGE: blue-native polyacrylamide gel electrophoresis
CBB: coomassie brilliant blue
CEF: cyclic electron flow around photosystem I
CO_2_: carbon dioxide;]
ECS: electrochromic shift
ECS_*t*_: total electrochromic shift from light to dark
FR: far red
*g*_H_^+^: transthylakoid proton conductivity
IR: inverted repeat
LEF: linear electron flow
NADPH: nicotinamide adenine dinucleotide phosphate
NPQ: nonphotochemical quenching
PCR: polymerase chain reaction
*pmf*: protonmotive force
*pmf*_LEF_: protonmotive force generated from linear electron flow
PQ: plastoquinone
PQH_2_: plastoquinol
PSI: photosystem I
PSII: photosystem II
q_E_: energy-dependent quenching
SDS-PAGE: sodium dodecyl sulfate polyacrylamide gel electrophoresis
WT: wild type
τ_ECS_: lifetime of the electrochromic shift decay during a dark interval
ΔpH: fraction of *pmf* stored as pH
ϕ_II_: quantum yield of photosystem II
*v*_H_^+^: trans-thylakoid proton flux

## Acknowledgements

We thank Dr. B. Sabater (Universidad de Alcalá) for generous donation of the *N. tabacum ndhF* mutant seeds Michael Tillich (MPI-MP) for generous donation of the *N. tabacum ndhA* mutant seeds, Luisa Heinig (MPI-MP) for help with tissue culture and transformation, and the MPI-MP GreenTeam and Sandra Stegmann for plant cultivation. We also thank Drs. David Kramer, Nicholas Fisher (Michigan State University), and Daniel Karcher (MPI-MP) for their helpful discussions. This work was supported by a grant from the Deutsche Forschungsgemeinschaft to R.B. (BO 1482/17-2; FOR 2092) and by the Max Planck Society.

## Author contributions

**Deserah D. Strand:** Conceptualization, Methodology, Writing; **Stephanie Ruf**: Methodology; **Omar A. Sandoval Ibanez:** Conceptualization, Methodology; **Ralph Bock:** Conceptualization, Methodology, Writing.

